# Neural ensemble activity depends on stimulus type in mouse primary visual cortex

**DOI:** 10.1101/708636

**Authors:** Marie Tolkiehn, Simon R. Schultz

## Abstract

Early cortical processing of visual information has long been investigated by describing the response properties such as receptive fields or orientation selectivity of individual neurons to moving gratings. However, thanks to recent technological advances, it has been become easier to record from larger neuronal populations which allow us to analyse the population responses to probe visual information processing at the population level. In the end, it is unlikely that sensory processing is a single-neuron effort but that of an entire population. Here we show how different stimulus types evoke distinct binary activity patterns (words) of simultaneous events on different sites in the anaesthetised mouse. Spontaneous activity and natural scenes indicated lower word distribution divergences than each to drifting gratings. Accounting for firing rate differences, spontaneous activity was linked to more unique patterns than stimulus-driven responses. Multidimensional scaling conveyed that pattern probability distributions clustered for spatial frequencies but not for directions. Further, drifting gratings modulated the Shannon entropy estimated on spatial patterns in a similar fashion as classical directional and spatial frequency tuning functions of neurons. This was supported by a distinct sublinear relationship between Shannon entropy and mean population firing rate.

## Introduction

Given the large number of neurons in the cortex, it is unlikely that visual processing is achieved on a single neuron level. Thus, to understand visual information processing in primary visual cortex it is important to examine neural ensembles and the signalling strategies between different groups of neurons. Population activity studies showed that different ensembles can be linked to certain stimuli and prefer to fire together^1–3^, that they can be inherently linked^4^, that the population activity is in fact coupled to the overall excitability (up or down states)^5^, and less influenced by visual stimulation itself^6–8^, or related to the strength of the stimulation^9^. Progress in experimental recording techniques allowed researchers to increase the number of simultaneously recorded neurons^10^. However, sophisticated analysis techniques are required to analyse such large amounts of neurons and their interactions. One measure that lends itself to compare neural data under different conditions is the simultaneous firing configuration^11,12^ of discretised and binarised individual spike times of each channel or neuron, henceforth termed patterns. These patterns may have different probabilities of occurring depending on what process or context generated them. For example, a pattern reliably evoked under visual stimulation of e.g. a drifting grating may be unlikely to be observed under grey screen (representing lack of stimulation) or a different type of stimulus^4^. It has been shown that activity of populations, discriminability and spiking reliability are contingent on both sensory inputs and internal states^3,13^. Spontaneous activity (SA), neural events present during quiescent wakefulness and passive attention, has been hypothesised to play a role in memory recall and consolidation, and in gating of sensory inputs^14,15^. SA can be similar in shape and appearance (e.g. synchronous events, correlated activities) to sensory responses^16–19^. This implies that the same networks underlie cell activity in both SA and driven contexts, or: There should be no difference between intrinsic probability of observing a particular pattern and under visual stimulation. Yet, other effects such as locomotion have been shown to enhance stimulus encoding and thus influence pattern probability^20^. With these differences in pattern occurrence, comparing pattern distributions under varying stimulus conditions is non-trivial particularly for large state spaces.

In this study, we investigate how different stimulus conditions and ongoing activity affect population responses in V1 of anaesthetised mice. We hypothesise that artificial stimuli such as moving gratings drive a distinct subset of pattern space occurring in spontaneous activity, and that stimuli mimicking those observed naturally, such as high spatial frequency features of natural scenes appear more similar to ongoing activity. Different mechanisms or stimulus components can be responsible for pattern generation^4,12^, which may lead to distinct subsets of the state space being activated under certain stimulus conditions. For example,^19^ showed that neurons were part of ensembles that fire in precise spatiotemporal sequences both under stimulation and absence of sensory inputs. They further showed it was possible to use these intrinsic activities to predict future temporal sequences under stimulation. While this may be an extreme case, variations of overlapping subsets^21^ or sampling biasses disguising the origin of differing populations may exacerbate the quantification process. Comparing individual pattern probabilities quickly becomes infeasible given limited sample sizes and many unobserved states^22–24^. That is why it is important to identify statistical derivatives and to pursue more holistic approaches, particularly those that can handle such problems^25^.

Here, we applied state-of-the-art data analysis techniques with particular focus on information-theoretic approaches such as Shannon entropy and Mutual Information (MI) on high-pass filtered thresholded signals. Information content was quantified for spatial patterns and contrasted with that contained in the population rate (i.e. summed activity across the neurons or channels, discarding spatial information). With the help of Shannon entropies, we computed the Jensen-Shannon Divergence (JSD) between pattern distributions and used it to examine how similar the distributions were during varying stimulus conditions, probing if the traversed pattern spaces were stereotyped for certain stimuli or stable under all stimulus and quiescent conditions. Our analysis showed that stimulus responses were highly reliable during grating stimulation and variable during stimulation through natural movies. Spontaneous activity evoked more unique patterns than stimulus-driven activity after accounting for firing rate differences. Further, pattern probabilities during spontaneous activity and natural movie stimulation appeared more similar than either to moving gratings. Analysis of pattern probability distributions revealed that spatial frequencies (SF) but not orientations appeared clustered on a subspace. This is an important step in trying to unveil how the brain encodes and transmits stimulus-dependent information.

## Results

We probed neural activity via 4-shank translaminar in-vivo electrophysiology (Neuronexus A4×8 silicon microelectrode, site spacing 100 *μ*m, shank spacing 200 *μ*m) in left hemisphere V1 of the isoflurane-anaesthetised mouse under visual stimulation of the right eye with monocular full-field drifting gratings, natural scene movies and SA (grey screen mean luminance). In particular, we investigated the binary firing vectors (pattern distributions) and population firing rates and applied information-theoretic approaches such as Shannon entropy and JSD to evaluate the similarities of SA and evoked responses. Analysis was based on simultaneous firing activities on recording sites of varying numbers of adjacent linear shanks (two or four), permitting information-theoretic analyses at different state space sizes to examine low-level visual processing and neural encoding. Wherever applicable, *stimulus type* or *stimulus condition* refer to the sets of data involving moving gratings (mG), spontaneous activity 1 (S1), natural scenes (nat), S2, temporal frequencies (TF) and S3.

### Reliable stimulus responses during gratings and high response variability during natural movies

Drifting gratings from the protocol presented in Fig. 1 (A) reliably evoked neural responses (Fig. 1 (B, C)). A high fraction of channels demonstrated strongly driven responses to visual stimuli, with stimulus-dependent response types. Moving gratings increased the firing rates (FR) with reliable on and offset responses across the population (Fig. 1 (B)), whereas natural movies induced spikes at a higher trial-to-trial variability and high population synchrony (C). The most reliable population response to the excerpt of the 30 s movie emerged as the onset response at t=0 s.

**Figure 1.**
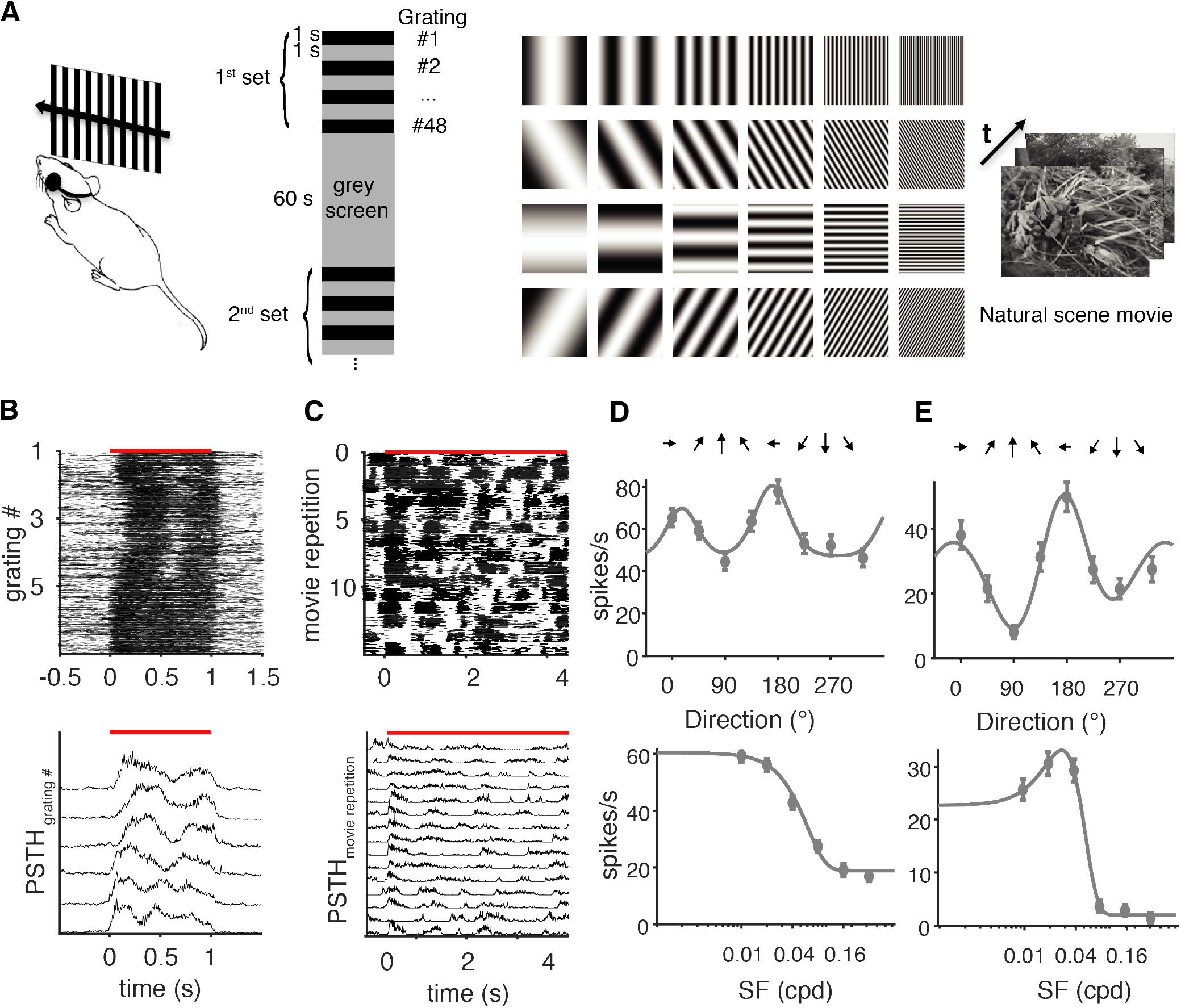
Grating stimuli and stimulation protocol. Response patterns vary between moving gratings and natural movies. (A) Left: Moving gratings are presented on a monitor covering approximately 60°x75° of the visual field of the right eye, while the contralateral eye is covered to avoid confounding effects from inputs from the contralateral eye; and in-vivo electrophysiological recordings were made from left hemisphere V1. Middle: Stimulus presentation (black) with interleaved spontaneous activity (grey), in pseudorandom order. Moving gratings are presented with 1 s *on* and 1 s *off* time, interleaved with 60 s grey screen, SA. Right: From left to right increasing spatial frequencies from 0.01 to 0.32 cpd, and from top to bottom increasing orientations at 45 ° steps, from 0/180 to 135/325. (D) Top: Example raster plots on 32 grouped channels (population response) to moving gratings at 6 directions (0-225° in 45° steps) at (0.01 cpd) for 20 repetitions, and to 15 repetitions of the first 4 seconds of the natural movie (E) illustrate visual responsiveness and spiking reliability across repetitions in the same animal. Each line highlights a multiunit spike event. (D,E) Bottom: Normalised PSTH for the same data, respectively. Red line indicates stimulus presentation period. (F, G) Direction (top) and SF (bottom) tuning functions for example sites from 2 mice show strong FR modulations. Direction and SF tuning curves appear stereotyped across mice. Top and bottom show responses from the same site. For direction tuning, FR were baseline-subtracted, averaged across the two lowest SF and repetitions. Baseline-subtracted FR of SFs were averaged over 8 directions and stimulus repetitions. Note bottom uses logarithmic x-axis scale, and note different y-axes scales. Error bars denote SEM.

The differences in the temporal structure of the Peri-Stimulus Time Histogram (PSTH) in Fig. 1 (B), bottom, suggested substantial tuning of the sites, both individually and population-wide. Figs. 1 (D) and (E) present two examples of strongly directionally and spatially tuned sites of two animals, whose shapes are representative across mice. This modulation was present at different average FRs. The tuning functions of these examples revealed a global minimum at 90°, and a maximum at 180°, where the direction 180° appeared to drive the activity slightly more than its collinear direction at 0°. Spatial tuning functions indicated drastic decreases in FRs for SFs exceeding 0.04 cycles per degree (cpd) (D, E, bottom). Sites displayed either low-pass (D) or bandpass properties (E), where low-pass was defined by a monotone decreasing shape, and bandpass behaviour by a rise-fall shape in the amplitude-frequency plot.

### Drifting gratings elicit more unique patterns than natural movies or spontaneous activity

After discretising the neural signals by binning the multi-unit spike trains in 5 ms bins, we binarised the signal as illustrated in Fig. 2 (A). This resulted in binary vectors of size *n* × 1 of the spike response over n sites for each time t, which we call (spatial) patterns, such that there is a total of *A* = 2^*n*^ possible patterns (words). The frequency of each observed pattern was then quantified (histogrammed) creating the pattern distribution.

**Figure 2.**
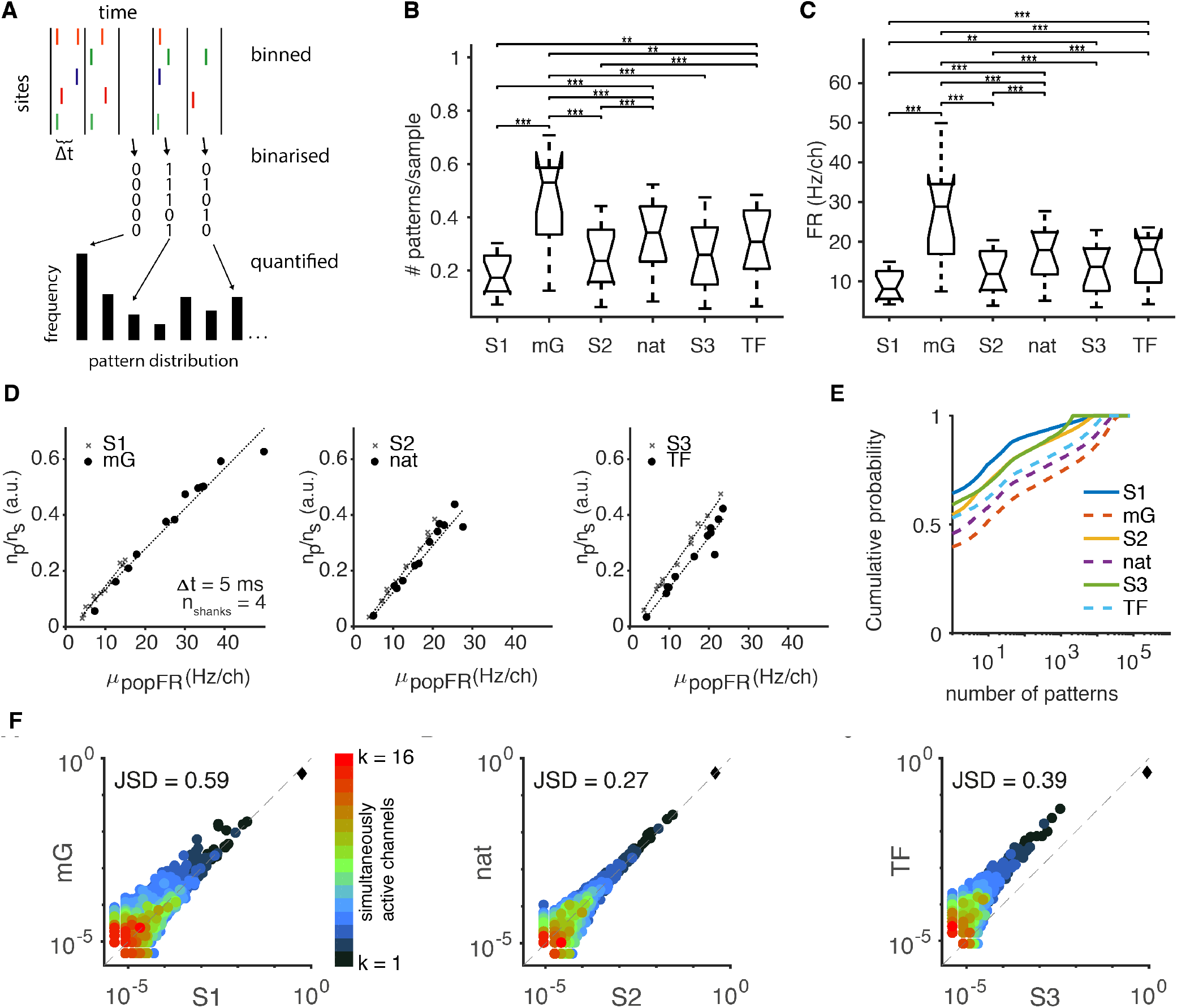
Drifting gratings elicit more unique patterns than natural movies or spontaneous activity. (A) High-pass filtered, thresholded events (spikes) of each site were binned (top) and binarised (middle), resulting in binary spatial patterns (or words), whose occurrences were quantified resulting in pattern distributions. (B) Number of unique patterns per sample (while accounting for varying recording lengths by subsampling) in box-and-whisker plots over all 32×1 patterns for the different stimulus conditions in 12 mice. (C) Mean population FR/channel varies with stimulus conditions (in 12 mice). In (B,C): ** indicates p<0.01 and *** p<0.001, Wilcoxon signed ranks, Bonferroni corrected. (D) depicts the number of patterns (*n*_p_) per number of samples (*ns*) against the mean population FR for stimulus conditions mG, nat and TF and their SA for 32-bit patterns at 5 ms bins. Dotted lines indicate linear fits. Each point is one mouse. (E) shows the (subsampled and averaged) cumulative pattern probability as a function of the number of patterns for each stimulus type in 32-bit patterns for an example mouse. Stimuli are shown in dashed lines, SA in continuous. (F) Empirical pattern probabilities vary under different stimulus conditions for 16-bit spatial patterns. The diagonal line indicates identity. Each dot represents one unique pattern (probability). Shades indicate number of simultaneously active channels (0-16). Diamond shape represents zero-pattern. Jensen Shannon Divergence between the distributions is indicated in top left corner. Left: Pattern probabilities during mG against S1. Middle: nat and S2. Right: TF and S3.

The number of unique patterns varied among stimulus types (Fig. 2 (B)). Attempting to account for differences in recording lengths, we subsampled from all categories 30 times, calculated the average number of patterns and divided it by sample length (16 · 10^3^ samples for all stimulus conditions). Spontaneous activity generally traverses a smaller set of unique patterns than evoked activities. Spontaneous activity (S1, S2 and S3) generated only approximately half the number of unique patterns of those present during stimulus presentation. The median number of patterns during mG amounted to almost twice those of its associated SA, with similar yet less pronounced differences for nat and its associated SA, S2. Another decrease in total number of patterns was visible for TF and S3, attributable to reduced recording length. A Friedman test (non-parametric version of repeated-measure ANOVA) yielded a highly significant influence of stimulus type on the number of uniquely evoked patterns/sample (p < 1.0e-299, *α* = 0.05, df = 5, *χ*^2^ = 1621.62, n = 360, achieved by 30 times resampling). According to Fig. 2 (B), the median number of unique patterns per sample lies approximately around 0.25 for most of the stimulation types, and is significantly higher during mG with a median over 0.5 patterns / sample, and a decreased value at S1 with a median just under 0.2. Stimulus presentation generally increased the number to 0.3. Moreover, it is apparent that the number of patterns per sample during S3 was approximately the same (non-significant) as during stimulus presentation TF. We explored the influence of stimulus types on FRs in Fig. 2 (C). This is expressed as mean population FR per channel to obtain the same numbers of data points as in (B). The two measures form concordant pairs wrt. stimulus types. SA appeared generally lowest, mG arose with highest median FR, and nat and TF exceed their respective SA. In accordance with results of (B), stimulus type significantly influenced mean population FR, *μ_popFR_* (p<1.0e-299, df= 5, *χ*^2^ = 1799.09, Friedman test, n = 360, achieved by 30 times resampling). Multiple comparisons of the medians pinpointed the highly significant difference in medians among mG and all other stimulus types (all p<0.001 Wilcoxon signed ranks, Bonferroni corrected). Further, in contrast to (B), in (C) S1 and S3 also differed significantly at p<0.01 (Wilcoxon signed ranks, Bonferroni corrected).

Following the concordance of medians in Fig. 2 (B,C), Fig. 2 (D) explores this further. The number of patterns per sample is highly positively correlated with *μ_popFR_* for all stimulus types and SAs (p<10^−4^, Pearson correlation coefficient >0.96 for each condition, mean correlation coefficient 0.98 ± 0.005 sem). Each group was fitted with a linear function of the type *y* = *mx* + *b* and obtained a minimum *R*^2^ of 0.88 (mean *R*^2^ = 0.94 ± 0.02 sem). Function fits were all very similar across stimulus conditions, and the function parameters are presented in Tab. 1. Y-intercept was slightly negative throughout, and the slopes ranged from 0.01 and 0.02 (a.u.), with all of the SA slopes exceeding the evoked ones. Evaluating regression results across all 6 stimulus conditions (all-to-all), the six slopes indicated no difference (p>0.05, F-statistic 0.01, df = 5, ANCOVA). Pooling each stimulus with the SA recorded in close temporal coherence allowed the comparison of slopes between recording windows. ANCOVA indicated a significant difference of slopes at p=1.9e-03 (F-statistic 252.60, df = 2, n: resampled 30 times). Comparing stimulus types (stimulation vs. spontaneous activity), the difference of slopes between grouped SAs and grouped stimuli resulted to be highly significant at p=1.0e-111 (F-statistic 568.52, df= 1, ANCOVA, n: resampled 30 times).

**Table 1.** Parameter values for linear fits depicted in Fig. 2 (D-F). Fitting functions are of the type *y* = *mx* + *b*. Individual slopes were not significantly different (p>0.05, F-statistic 0.01, df = 5 ANCOVA), slopes between grouped SA and grouped evoked conditions differ highly significantly (p=1.0e-111, F-statistic 568.52, df = 1 ANCOVA, repeated subsampling 30 times).

|  | intercept b (a.u.) | slope x (1/Hz) |
| --- | --- | --- |
| S1 | -0.0352 | 0.0181 |
| mG | -0.0093 | 0.0144 |
| S2 | -0.0429 | 0.0190 |
| nat | -0.0451 | 0.0168 |
| S3 | -0.0086 | 0.0206 |
| TF | -0.0415 | 0.0182 |
| Grouped SA | -0.0457 | 0.0211 |
| Grouped evoked | -0.0022 | 0.0147 |

Fig. 2 (E) illustrates the cumulative probabilities of 32-bit patterns as a function of number of patterns for an example mouse. Probabilities were repeatedly subsampled and averaged to account for different recording lengths. The y-intercept corresponds to the zero-pattern, which is visibly lower under stimulus presentation (dashed lines). Especially mG indicate many more activity patterns to account for the same probability mass as e.g. S1.

We compared pattern probabilities under all stimulus conditions to enquire into whether the pattern statistics varied with stimulus types, or if distinct neural ensembles were involved. Fig. 2 (F) displays scatter plots of pattern probabilities under different stimulus conditions in an example mouse, for 16-bit patterns on log-log axes. Colour map corresponds to number of simultaneously active channels (0-16, blue to red), with a black diamond indicating the zero-pattern, which was the most frequent one in all sets and animals. In all cases, patterns with a small number of simultaneously active channels exhibited high probabilities, and patterns with a large number of co-active sites had generally lower probabilities, accumulated in the lower left corners. As can be seen in Fig. 2 (A, left), the zero-pattern occurred at a slightly higher probability in S1 than during mG. Analogously, a vast subset of patterns occurring during S1 arose under mG-evoked activity with much higher probability, shifting away upwards from the identity line. This was particularly obvious for higher spike synchrony patterns, which clustered more above the identity line at low probability values. This was reflected well by the fairly large JSD between the two distributions, which amounted to 0.59 bits. The middle panel reflects the probability scatter between nat and its associated SA, S2. The zero-pattern appeared to be almost equally frequent in the two conditions, as suggested by the black diamond on the identity line. In addition, most pattern probabilities distributed axisymmetrically around the identity line for both low and high spike-count patterns, with increasing spread at low probabilities. The JSD was substantially lower than during gratings and their SA at 0.27 bits. Finally, the right panel compares TF and S3, which appear in shape similar to the middle panel with a tight spread shifted away from the identity line towards TF, with considerably higher pattern probabilities for the evoked case. The zero-pattern was again displaced from the identity line, and JSD amounted to 0.39 bits. These findings were qualitatively similar across mice, and generally, the JSD was highest between S1 and mG, and lowest between nat and S2.

### Neural ensemble spatial pattern entropy weakly resembles tuning functions

In the following, 16×1 binary words (patterns) were computed by taking 2 adjacent 8×1 shanks in each mouse, resulting in 24 non-overlapping experimental sets (i.e. 2 for each of the 12 mice), bins were assumed temporally independent. A subset of the channels was used because a reliable Shannon entropy estimate of 2^32^ patterns would have required many more samples.

Fig. 3 visualises pattern entropies (A) for different stimulus groups and types. The mean population FRs across the same set is shown in (B). Grating pattern entropies in (A) were estimated on the probability distributions over all directions at the lowest four SF, in order to condense information about directions, as was presented in Fig. 1 (D, E, bottom) that higher SFs evoked less distinguishable responses. To complement this, in the right panels of (A, B), entropies are estimated over the pattern distributions over all directions for each SF separately. Inspection of Fig. 3 (A, B) suggests that pattern entropies resemble mean population FRs across all stimuli and also seem to weakly approximate the tuning curves (cf. Fig. 1 (D,E)), with peaks around 180° and troughs around 90° across mice for the directions. Correspondingly, entropy over SFs roughly resembled the DoG tuning fits observed in Fig. 1 (D, E), with higher values at low SFs and lower values at high SFs. This grouping also allows enquiry into where the other stimulus types, S2, nat, S3 and TF approximately lie in relation to the tuning functions of Fig. 1 (D,E). As illustrated in Fig. 3 (A), S1 and S2 manifest the lowest entropies apart from TF, with nat displaying an entropy not much higher. In accordance with this, in (B) mean population FRs (*μ_popFR_* in Hz/channel) estimated over the same samples of (A) exhibit the same qualitative shapes across stimuli. *μ_popFR_* is low for spontaneous activities, shows a dip around 90°, and a peak at 180°. Similar to the entropies, *μ_popFR_* increases slightly again for nat movies.

**Figure 3.**
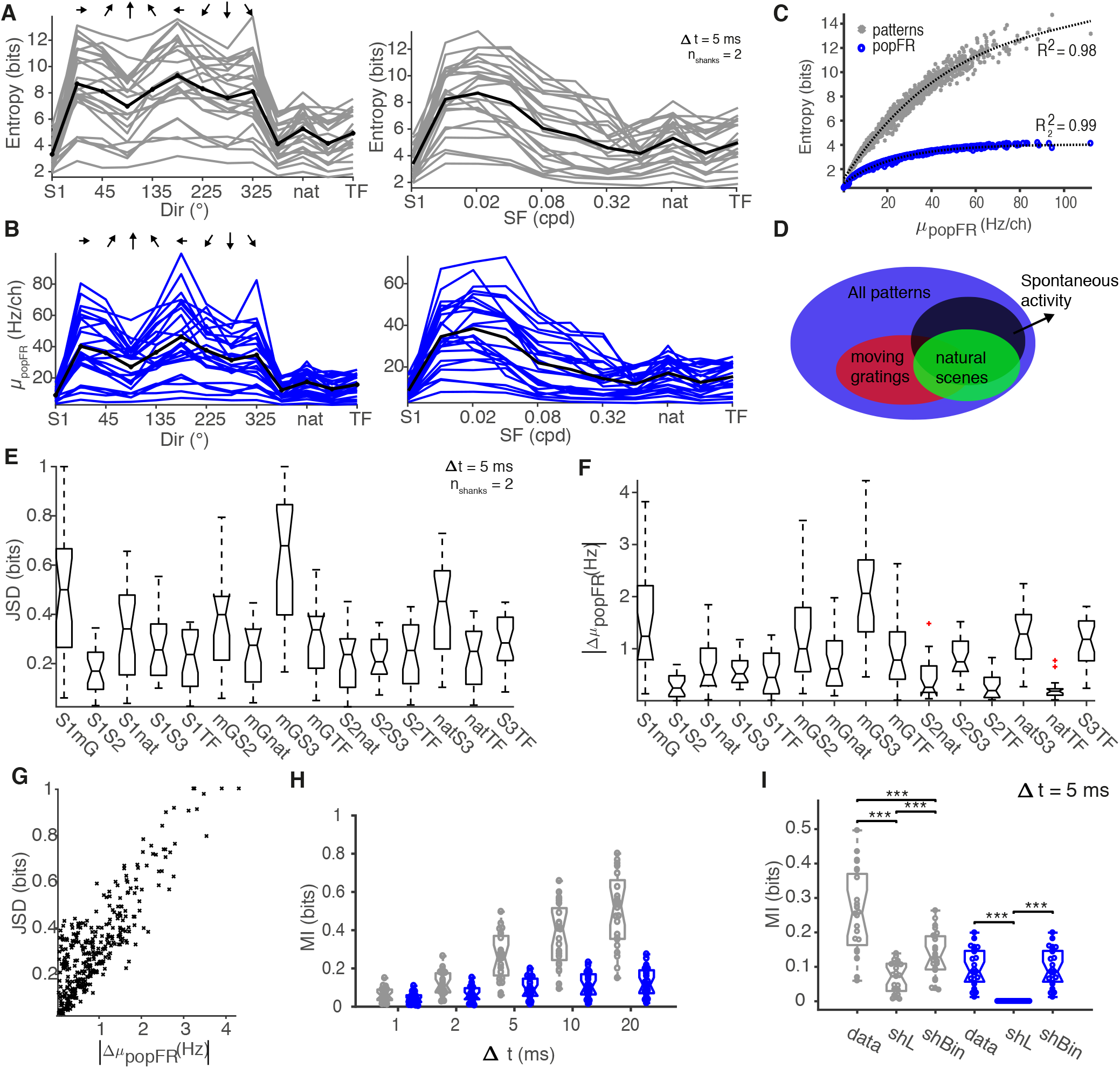
Shannon entropy and *μ_popFR_* follow a sublinear relationship. (A) Each line represents entropies computed over 16×1 binary patterns at 5 ms bins for each stimulus type of one of 24 experimental sets (n=12 mice). X-axis shows S1, moving gratings (left: 0-325° in 45° steps, right: 0.01-0.32 cpd), S2 and nat movies, S3 and TF. Left: Pattern entropy over directions, pooled over the four lowest SFs. Right: SF entropies, pooled over all eight directions. (B) same as in (A) but for mean population FR in Hz/channel computed over the same samples. (C) Entropies computed on patterns (grey) and population FR (blue) increase sublinearly with *μ_popFR_*. Each symbol corresponds to one of 62 (48 gratings, 10 TF, 1 nat, 3 SA) stimulus cases in 24 experiment sets. (D) Schematic of how stimuli occupy different subsets of pattern space. (E) Pairwise JSD distributions over stimulus presentation types show e.g. large divergences between S1 and mG and small divergences between e.g. S1 and S2. (F) *Aμ_popFR_* between the same conditions as in (F) in Hz/channel appear in concordance with divergences from (G). This is further illustrated in (F), which shows one ‘x’ per stimulus condition for all mice indicating a strong positive correlation between difference in FRs and JSD (r=0.85, p<0.0001, Pearson). (H) Mutual Information (MI) between spatial 16×1 patterns and 62 stimuli computed on patterns (grey) and population FR (blue) emphasises the superiority of patterns over population FR, particularly with increasing bin size. (I) MI between patterns and stimuli at 5 ms bins for patterns (grey) and population FR (blue), with shuffled labels (shL) and shuffled bins (shBin). *** indicates p<0.001, 2-sample sign test.

Entropy increases with average population FR in a positive sublinear relationship (C). This is not only true for pattern entropies, but also entropies estimated using population FRs alone (entropies calculated with the Pitman-Yor-Mixture (PYM) estimator on the population FR). Population FR entropy is much lower than pattern entropy (17 vs 2^16^ possible states for patterns, p<0.001, Wilcoxon signed ranks test), but both increase sublinearly with mean population FR. They are fitted with a cubic of the form *ax*^3^ + *bx*^2^ + *cx* + *d* at high *R*^2^ values of 0.98 and 0.99, respectively.

### Stimuli occupy different pattern spaces

This difference in entropies suggests that moving gratings occupy a different, larger pattern space than spontaneous activity or moving gratings, as illustrated in Fig. 3 (D). To quantify the similarity between pattern distributions, we computed the JSD between each of the possible combinations of stimulation sets.

It is difficult to disentangle influences by differences in FR from the disparity in pattern probability distribution induced by the different stimuli themselves. To investigate this further, Fig. 3 (E) depicts the distributions of JSD and (F) the distributions of the *difference* in mean population FR among stimulus conditions. There appears to be a relationship between JSD and the difference in *μ_popFR_* across stimulus conditions, Δ*μ_popFR_*. Fig. 3 (E) summarises the JSD of all stimulus presentation types, again illustrating the large divergences between e.g. mG and S1. The box plots indicate the median (central mark), the bottom and top edges the 25 and 75th percentile. Whiskers point to the most extreme data points, and outliers are illustrated as red crosses. Data points are classified as outliers if they exceed *q*3 + *w* * (*q*3 − *q*1) or less than *q* − *w* * (*q*3 − *q*1), where *w* stands for maximum whisker length, and *q*1 and *q*3 are the 25th and 75th percentiles. To compare the qualitative progress of JSD values to the differences in FR, Fig. 3 (F) discloses Δ*μ_popFR_* for the same presentation types in Hz/channel. Contrasting (E) and (F), a concordance in value progression is evident, which is further highlighted in (G) illustrating all JSD - FR pairs of all mice and stimulus conditions. Fig. 3 (G) indicates a positive correlation between divergences and difference in average population FR despite displaying a large variance particularly at low values (r=0.85, p<0.0001, Pearson, n=24×15).

### Mutual Information between neural activity and stimuli

Quantifying the information content of the spatial patterns and stimulus (total 62 stimuli) in terms of MI revealed that the spatial patterns contained 0.26 bits of information (median, cf. Fig. 3, (H)) at 5 ms bins. MI is maximal for empirical spatial patterns, in contrast to estimating MI with population FR, where the median amounted to 0.09 bits (at 5 ms bins). Increasing bin widths greatly increased MI estimates. Yet, larger bin sizes might positively bias the estimate through the decreased number of samples (thus negatively biassing entropy estimates). MI is highest when estimated using spatial patterns. Fig. 3 (I) illustrates MI computed for patterns and population FR at 5 ms bins, and how randomly shuffling stimulus identities (labels) for each sample, or randomly shuffling spatial patterns site-by-site for each bin reduces MI.

Shuffling stimulus labels greatly reduced MI over the unshuffled results (Fig. 3 (I)). This is particularly striking for MI calculated via population FR, where MI is reduced to nearly zero (small positive fluctuations) for shuffled labels (shL, p<0.001, 2-sample sign test). MI estimated via pattern entropies was greatly reduced for shuffled labels (p<0.001, 2-sample sign test), but indicated residual information possibly attributable to negative entropy biases. In addition, randomly shuffling bins within a sample (spatial shuffling, shBin) reduced the MI estimates greatly for patterns (p<0.001, 2-sample sign test). Trivially, spatial pattern shuffling does not affect population FR, as the sum across sites remains unchanged.

### Pattern probabilities evoked by natural movies and spontaneous activity diverge less than either moving gratings

Using pairwise JSD of stimulus presentation types, we can compare pattern distributions beyond number of patterns and quantify how dissimilar the distributions are. This may indicate that certain stimulus types traverse similar patterns with similar probabilities or that some patterns belong to a different distribution, if they do not or rarely occur in other stimulus types. Multidimensional Scaling (MDS) is a technique that takes in a distance matrix and transforms it into a lower dimensional space whilst seeking to preserve distances (between distributions in our case). Using pairwise JSD as the input distance matrix, we can visualise the pattern similarities in a simple graph illustrating how distributed the stimulus types are in relation to each other. This is delineated in Fig. 4, indicating where the pattern distributions lie with respect to each other in an artificial space.

**Figure 4.**
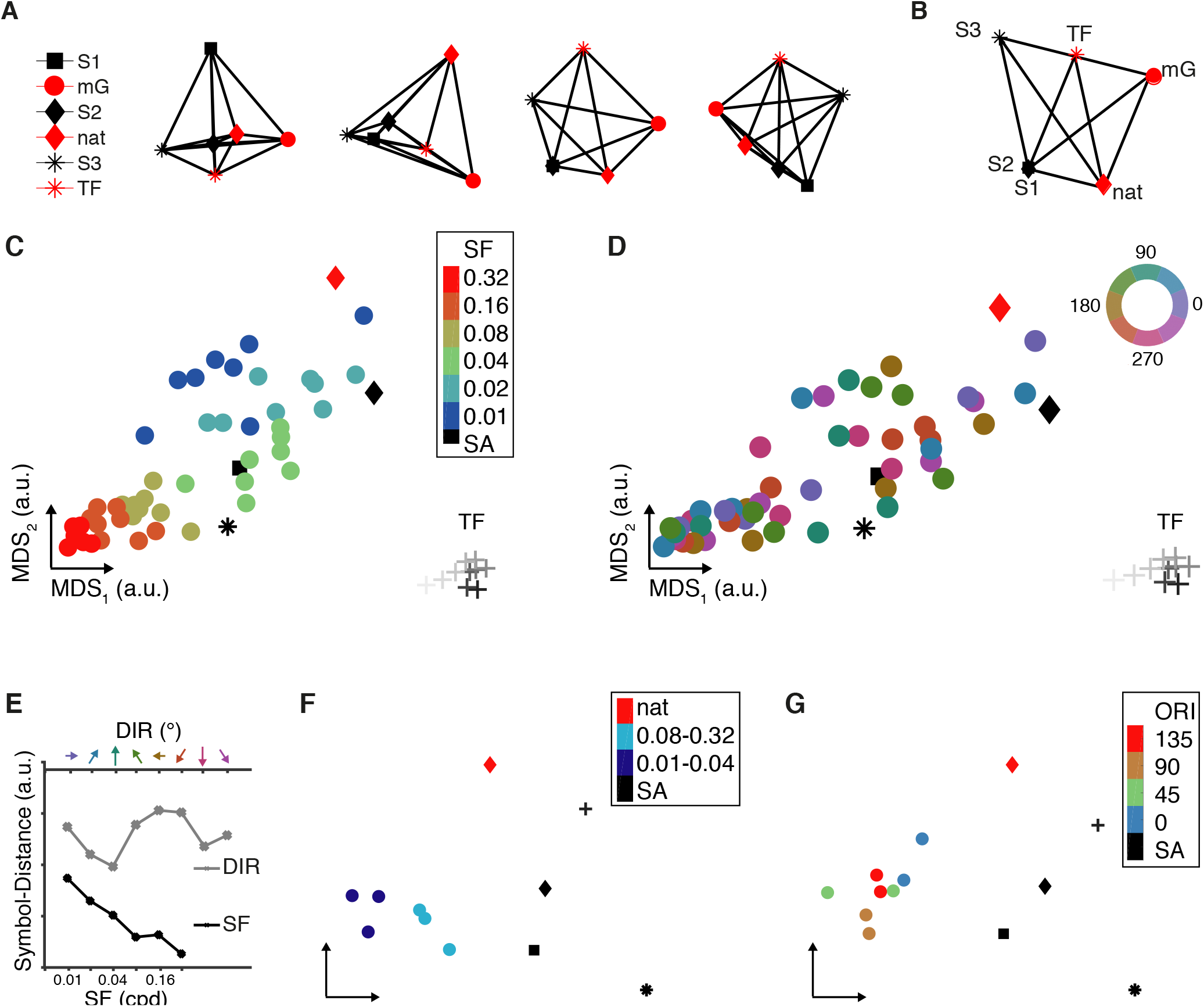
Pattern distributions conditioned on all stimuli appear clustered for spatial frequencies, and less structured for directions. Each symbol represents one stimulus type, colour indicates spontaneous activity (black) or stimulation (red). (A) Four examples of illustrating the low dimensional mapping using Multidimensional Scaling (MDS) on the JSD matrix. (B) shows the MDS graph on the average JSD over 12 mice. (C) MDS on the mouse-averaged JSD distance matrices of the pattern probability distributions unveils a map clustering in SFs, whereas DIR seem less globally structured in (D) with a circular colour map. In all subpanels, the natural movie sets (black and red diamond), are consistently at a distance from mG. Colours represent same SF or direction, respectively. Square indicates S1, diamonds S2 (black) and nat (red), plus-signs TF (grey scale) and S3 (black star). (E) Intra-class symbol distances decrease for SFs, but not for directions. NB: different x-axes. (F) Pooled directions at each SF cluster into high and low spatial frequencies. (G) DIRs pooled across SFs appear very similar to each other in contrast to S1, S2, S3 and nat and TF. Colour code represents same orientations, for clarity.

Fig. 4 (A) visualises the lower dimensional mapping (a.u.) for 4 example data sets (estimated on 16-bit patterns of 4 mice). Each symbol stands for one stimulus type, colour code indicates visual stimulation (red) and SA (black). Scrutiny of (A) evinces a close relation between nat and S2 in the centre of the graph (diamond shapes), while the remaining evoked types (mG and TF) appear in the vicinity and the spontaneous cases are located further away. The next example offers a different graph in that SA and evoked types are more distinctly clustered, with all SA in a tight neighbourhood, and evoked activities displaced, where TF is on a trajectory between SA towards mG, and nat appears in a different regime. The next mouse maps SA of S1 and S2 in the same spot, with the nearest evoked distribution again being nat. The fourth example displays a symmetric graph with the symmetry line at roughly 45° marking the difference between evoked and spontaneous distributions. The edge between nat and S2 is the shortest, with mG and S1 aligned to it on either side, and TF and S3 further removed, forming a trapezoidal shape. Graph (B) is the low-dimensional MDS mapping on the *mouse-averaged* JSD matrix. The mean JSD over all mice results in a short edge length between nat and SA in contrast to drifting gratings (both mG and TF), which are located further away.

### Spatial frequencies are clustered in pattern space

Using the same technique to build the JSD distance matrix across all stimuli (S1, 48 mG, S2 and nat, S3 and ten TFs, creating a 62×62 matrix) allows us to visualise the combinations of SF and directions in pattern space. In this case, divergences are estimated over unequal numbers of samples. Sample sizes for spontaneous activities and nat movies are unchanged from before, but grating sample sizes are now reduced to 20 repetitions of 200 5 ms bins (i.e. 20 000 samples each).

Fig. 4 (C) highlights the MDS map over the mouse-averaged JSD distance matrix. Each circular symbol represents one grating type (of defined SF-direction combination), and colour of circular symbols corresponds to their SF. Black symbols depict SA, diamonds the set of nat movie, the star S3 and plus signs TF. It is evident from the figure that the arrangement is not random, but that gratings, in particular spatial frequencies form clusters in the transformed space. Again, it is apparent that nat movie patterns seem to be displaced from gratings at the top right corner (red diamond). However, SA recorded interleaved with mG appears to be in close proximity to mid-frequency gratings (green, 0.04 cpd), and almost central in the graph. The circular symbols appear not only clustered but also to be following a path at increasing SF, from blue to red, whilst decreasing inter-symbol-distance or scatter as well. All TF are far removed from the remaining stimulus types in the lower right corner, also following an apparent order of increasing TF (black to pale grey). Fig. 4 (D) illustrates the same mapping with a different colour code, where each colour represents one direction on a circular colour map. The colour appearance is more scattered, with some local clusters of directions. Local structures emerged e.g. for 45/225° (green) in the centre of the graph, corresponding to the lowest three SF of (C). Direct comparison between (C) and (D) suggests that orientations at the same SF tend to appear in close proximity to each other (e.g. two circular symbols at 0.02 cpd in (C), moss green, appear in brown 90/270° in (D)). Fig. 4 (E) shows the average within-class distances between symbols to quantify their scatter. Individual symbol distance was calculated as the Euclidean distance between symbols. Intra-class distance for SF decreases at higher SFs. Intra-class distances are generally higher for directions and do not appear to follow a clear structure. Inter-class distances were high for SF (not shown), given their localised clusters. Average distance between directions was small, given the large overlap among groups.

This observation was reinforced when grouping the data for SF alone (pooling trials over all directions for each SF), which is portrayed in Fig. 4 (F). Here, each circular symbol represents one of six SF, and JSD is calculated over all directions at the same SF. Again, the figure suggests a difference between nat movies, mG and SA. S1 appears in the general SA area, further away from mG, or, the mGs form a cluster distant from the remaining presentation types. Between mGs, we can identify a sub-clustering of SFs in lower (blue) and higher (turquoise) SFs, with lower SFs further away from the centre of the graph, and thus further away from SA of the right side. The other visual stimulations, nat and TF emerge at the top right corner forming a distinct contrast to the SF clusters. Similarly, Fig. 4 (G) pools all SFs with the same direction and calculating the pairwise JSD between the remaining 8+2+3 stimulus types reveals directionally distinct clusters: nat movies, TF, S1, S2 and S3 appear far away from the directions in similar locations as in the previous panel. In this representation, some of the directions emerge close to their collinear direction.

## Discussion and conclusions

This study applied information-theoretic approaches such as Shannon entropy and MI to binary pattern responses, to investigate how neural population activity was affected by stimulus types in V1 of anaesthetised mice. We examined how neurons encode and process stimulus-dependent information and how and if it differed in the absence of stimulation.

We found that the number of unique patterns in each experimental condition depended on the stimulus type, which was tightly linked to mean population firing rate, which failed to completely explain it. While S1 and S2 differed significantly in their patterns per sample, their FR were not found to differ significantly, suggesting a higher pattern yield in S2 that cannot be explained by FR alone. S3 was notable in its homogeneity of response patterns, demonstrating that natural movies (which preceded the S3 period) might evoke periods of stereotyped activity after cessation. One drawback of normalising by sample size (duration) was that the short presentation time of S3 may bias the results as it could artificially increase the ratio of patterns / sample if there was an actual minimum number of total patterns in that regime. Balanced recording lengths would have eliminated this concern. In line with increased unique patterns per sample, *μ_popFR_* was highest during mG, and we identified a strong positive correlation between number of unique patterns per sample and *μ_popFR_*. Yet, overall, spontaneous activity evoked more unique patterns than stimulus-driven activity when accounting for firing rates. Grouping SA and evoked responses resulted in highly significant differences between slopes (p<10^−6^, F-statistic 29.87, ANCOVA). In addition, the slope computed over the aggregated SA exceeded that of grouped evoked slope by approximately 40%. This implies that the pattern variety during SA is larger than in evoked activities, when accounting for the smaller population FR. Also, patterns during evoked activity appear more reliable or stereotyped as has been reported elsewhere^26, 27^, even to the degree that stimuli reduce the dimensionality of activity in line with findings by^28^.

We further found that entropies resembled spatial and directional tuning profiles of individual sites. Thus, tuning properties were visible at both single-site and population level and entropy properties appeared to be fairly stable across locations and mice (apart from scaling differences), suggesting a similar processing strategy. We demonstrated a strong positive relationship between entropy and *μ_popFR_* for both entropies estimated via spatial patterns and population FR. The fit between entropies and *μ_popFR_* indicated a plateau at about 4 bits (theoretical maximum is log_2_(17) = 4.0875 bits) for population FR-entropies, while entropy estimated via patterns attained higher values tending to its theoretical maximum of 16 bits. A sublinear relationship between entropy and *μ_popFR_* may reflect the sparsity of neural data. At low population FR, only few sites are active, and thus, only few patterns are observed, resulting in low entropy. At higher population FRs, more patterns can be traversed, resulting in higher entropy, particularly under the assumption of each site firing with equal probability. Equally, at higher population FR, the number of simultaneously active channels increases. This must reach a plateau once a certain FR is passed, and then decrease again. The conditional entropy of a response *R* given the number *n* of simultaneously active channels *H*(*R*|*n*) is Bernoulli distributed for independent channels. A toy example proves for a 2^16^ pattern, that 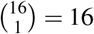 and 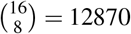, representing the number of possible combinations of patterns with one and eight simultaneously active sites. Picking the maximum *μ_popFR_* of approximately 100 Hz/ch (from Fig. 3 (C)), we can estimate how many channels were active on average, per bin. Given 100 Hz/ch was estimated over bins of size Δ*t* = 5*ms*. This means 200 bins of size 5 ms contained 0.5 spikes per bin and channel. FR were averaged across 16 channels, leading to averagely 8 sites being active simultaneously. Having 8 channels simultaneously active results in 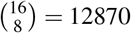 possible states, amounting to *H*(*R*|8) = 13.65 bits, which is roughly given in (E). This relationship must decrease at high *μ_popFR_*. For example, at 200 Hz/ch, each channel requires to be active simultaneously (the all-ones pattern), and entropy of only one state is zero.

In conjunction with this, MI between stimulus and response estimated via population FR obtained values around 0.1 bits, whereas MI computed using patterns attained values more than twice as high, largely increasing at higher bin widths, emphasising the importance of spatial representation. However, larger bin widths may positively bias the estimate since the numbers of samples decrease (c.f. negative bias from entropy), which may affect the pattern-derived entropies more than the population FR one. Shuffling labels reduced this information to nearly zero for population FR (p<0.001, 2-sample sign test), and resulted in a small positive bias for MI calculated via patterns. Spatially shuffling bins in the patterns significantly (p<0.001, 2-sample sign test) reduced the MI estimate, again supporting the importance of spatial spiking configuration. Differences in entropy estimates between shBin and population FR (both features without the spatial structure) may be attributable to the different entropy estimators used.

In addition, we found evidence that natural movies may be processed differently from moving gratings. Individual pattern probabilities during SA (grey screen) and nat movies appeared to be fairly similar, both visually and in terms of the JSD, whilst mG seemed to modulate pattern probabilities differently. The experimental design allowed for comparing SA recorded at different times during the experiment, which enabled us to account for different brain states or network excitability levels^5,8,13^ Particularly nat movies and S2 recorded in close temporal coherence (interleaved between stimulus repetitions) revealed a low divergence of pattern distributions with values comparable to divergences between SA recorded at early and late experimental phases. It was argued by Luczak^16^ that responses during SA may delineate the set of cortical responses, and the role of SA on sensory processing^14,29^. The fact that TF and S3 appeared furthest away in e.g. Fig. 4 may be linked to the timing in the experiment (later in the general anaesthesia) plus the difference in recording lengths, which may influence minimum possible probability. The former may reflect different brain states, a change in excitability or gain, or that these stimuli are processed differently, with nat movies being closer to the default firing states. Natural movies with their high SFs and large amount of spatial detail drive the neurons at a highly fluctuating pace, whereas mG can evoke responses that are more sustained during the presentation period because of their high contrast and generally large spatial edges. Yet, non-optimal gratings, which have similar average FRs as nat movie-evoked responses still elicited very different patterns, suggesting it was not FR differences alone. Berkes presented similar findings in ferrets^18^, which have structured orientation maps like cats, where particularly in adult ferrets mGs were most dissimilar to SA and nat movies. They postulated that a statistical model that is optimally adapted to a stimulus ensemble must have had prior experience to match the occurrence frequency and investigated the difference in pattern distributions (with a symmetrised version of the KLD) over the developmental period. Miller^4^ also argued for patterns observed during spontaneous activity to be linked to ensembles activated and formed during prior stimulus exposure. Our study ties in nicely here and confirms Berkes’s observation for adult anaesthetised mice.

Okun^30^ argued that word distributions differed between *cortical states,* where brain state was estimated as the coefficient of variation as derived in^31^. In their paper (on rats and cats primary visual and auditory cortex), brain state was the main factor for pattern similarity, more so than the presented stimulus (or lack thereof) and all correlations were subject to population rate dynamics. Unfortunately, their model was a poor fit if strong correlations between subgroups of neurons existed that could not be explained by population FR dynamics. This sparked lively discussions between the groups^32,33^. However, the present study confirms that the pattern distribution during SA is, also in mice, more similar to nat movies than to gratings, as was shown in ferrets^18^, rats and cats^30^. Further, our findings do not contradict Okun’s, that population rate dynamics may play an important role, as we recorded nat movies interleaved with SA in an attempt of matching or accounting for brain states throughout the recording. In a similar investigation in behaving monkeys Tan^34^ observed that visual stimulation shifts cortical state to the asynchronous state, which could be reflected by higher entropies we observed during visual stimulation.

Investigation of pattern distributions via JSD and MDS revealed that SFs appeared clustered in the MDS-transformed pattern space. SA and nat movies emerged at the periphery of pattern space, whereas SFs tiled the majority of it. Low SFs were clustered while retaining a high spread, and traversed in an orderly fashion to more densely packed patterns of higher SFs. The increase in pattern similarity at high SFs may reflect the detectability of the grating. High SFs may be more difficult for the mouse to detect, and thus evoked patterns may be represented similarly, and therefore, cluster. The larger spread for low SFs on the other hand may correspond to the differences in modulations by the directions. This directional difference may be washed out at higher SFs, resulting in less diverse pattern distributions. This may suggest that SF were the main driver for pattern space in the subcategory of mGs, and that this clustering can be subdivided at low SF to achieve a secondary tiling corresponding to directions of a grating.

Dissimilarities in pattern distributions between stimuli may be attributable to different FRs induced by the choice of stimuli (optimal gratings evoke high FRs whereas nat movies may induce on average lower FRs with instantaneously high peaks). Spontaneous and nat movie evoked activity displayed a lower JSD than both/each to artificial gratings. If neurons were more driven by (optimal) artificial stimuli, one might argue that this could automatically change the pattern distribution. However, non-optimal gratings with FRs similar to nat movies still appeared to elicit different pattern statistics. Further, Fig. 3 (E, F) showed that stimulus type pairings that differed less than 1 Hz achieved divergences en par with those with Δ*μ_popFR_* around 1 Hz. For instance, *natTF* indicated the minimum difference in FR but the median JSD amounted to 0.25 bits, similar to e.g. *mGnat*. Thus, mean firing rates do not completely account for the pattern distribution divergences. One way to check individual FR influences could be to create surrogate data (e.g. homogeneous independent Poisson that match the individual site firing statistics, which is the maximum entropy solution given FR constraints^35^). This would use the mean FR for each stimulus, but will not reflect population FRs. Alternatively, surrogate data that match only the *μ_popFR_* for each condition could be used to check if this dissimilarity is indeed dependent on population FRs induced by the stimuli.

In addition, it has been suggested^30,32^ that instead of pairwise correlations between spike trains, population FR fluctuations may be responsible for the different pattern distributions and that considering population rate dynamics over simply mean FR was required to account for FR statistics^36^ and that fluctuations in ongoing activity play an important role in population FR^13^. There has been considerable research into this with different models^13,24,30,36,37^. Population dynamics, and population FR differences between the various stimulus regimes could be implicated in the observed divergences.^37^ investigated the relationship between one neuron and the population. They coined the terms choristers and soloists for neurons that showed a tendency to be highly linked to population FR and independent thereof, respectively. Choristers appeared to be affected by sensory stimulation, while soloists are much less influenced by the ensemble or network activity. They further suggested that the coupling of neurons to the population FR explained pairwise correlations. Both soloists and choristers are reflected in Δ*μ_popFR_*, and thus, the choristers’ robustness in sensory stimulation^37^ could partially explain the consistency in our results.

## Methods

### Surgeries and animal preparation

We used 4×8-shank translaminar linear Neuronexus silicon microelectrodes (A4×8-5mm-100-200-177) to record neural activity from different layers and columns of left hemisphere mouse V1 under visual stimulation of the right eye. Mice were group-housed, kept in a reversed 12 hour dark/light cycle and recordings were performed during the early dark phase.

#### Anaesthetics and drugs

12 female young adult C57BL/6 mice (mean age 2.2 months) were sedated intraperitoneally with chlorprothixene (0.5 mg/kg, Sigma-Aldrich, UK) and anaesthetised with isoflurane (2% for induction, 1 – 2% for surgery, 1% for electrophysiology in 1.2% O_2_, Harvard Apparatus, UK). A vaporizer controlled the isoflurane concentration and a scavenger retrieved superfluous anaesthetic. To avoid tracheal secretions and to maintain clear airways, we injected 0.3 mg/kg atropine sulphate (Animalcare, UK) subcutaneously. To prevent oedema, we administered 2 mg/kg dexamethasone (Organon, UK) subcutaneously. We controlled the depth of anaesthesia continually by checking the pedal-withdrawal reflex.

#### Surgical procedures

After anaesthesia induction, the animal was moved onto a feedback-regulated heating pad. The body temperature was measured with a rectal thermometer and maintained at 37.1 ±0.5°C. To avoid corneal dehydration whilst maintaining clear optical transmission we applied eye ointment (silicone oil, Sigma-Aldrich), and covered it from microscope light. Head position was secured using ear bars, a custom-built nose cone and an incisor adapter. Our target location was the monocular region of primary visual cortex (V1m) at −3.55 mm posterior Bregma and 2.5 mm lateral from the midline^38^. We identified our target area by first stereotactically measuring the distance between Bregma and Lambda. This distance was used to correct the target location with *x_LOC_* = −3.55 mm* z[mm]/4.2 mm and *y_LOC_* = 2.5*z[mm]/4.2 mm, with z the individual distance between Bregma and Lambda, and 4.2 mm being the average literature distance between them^38^. The location for the ground screw over the contralateral cerebellum was identified on the interparietal bone near the lambdoid suture by applying PBS to avoid superficial blood vessels. We drilled a small craniotomy (1 mm diameter) for the ground screw with a hand-held dental drill (Osada Success 40, 0.5 mm drill bit). After removing the dura with a small needle (27G), the previously prepared ground screw-socket complex (Precision Technology Supplies, M1.0×2.0 Slot Cheese Machine Screw DIN84 A2 St/St and socket connector, MILL MAX, 851-43-050-10-001000 connector, sip socket) was inserted into the craniotomy. An elastic ring from a syringe tip or PVC tube, glued onto the target area with super glue (Henkel Loctite) served as a well over the craniotomy site. We secured the ground screw and well in place with dental cement (Kemdent Simplex Rapid®, cold cure acrylic). We also used the dental cement to cover the exposed skull and to form a head plate, joining the skull with a custom horizontal metal bar affixed to the frame. Once the dental cement set, ear bars were removed. For the 2-3 mm diameter craniotomy inside the well, we thinned the skull at the outer part of the ROI, and once the bone was thin enough to form cracks, we applied PBS, easing the bone fragment removal. Doing this facilitated the separation of skull and dura, minimizing potential damage due to skull-dura adhesion. The skull fragment was lifted slightly off the brain with a bent needle as a hook, and removed with fine forceps (Dumont #5). To remove the dura, we made a small incision lateral to the recording site, and retracted it with fine forceps #5. The mouse was then moved into recording position. The electrophysiology rig, modified from the set-up designed and used by^39^, was built on an air-pressure stabilised optical table with aluminium plates on four sides to form a Faraday cage, minimizing electromagnetic noise artefacts and scattered light. The front part of the rig was covered with a conductive fabric curtain (Wavetame, UK), drawn during recordings, keeping light and noise away from the recording sites.

#### Stimuli and data acquisition

Our stimuli comprised a set of pseudorandomly presented sinusoidal drifting gratings at full contrast, 20 repetitions each, of 6 different SFs at 8 directions (1 s stimulus-ON time, with 1 s pre-stimulus time) at a constant 1. 6 Hz temporal frequency, interleaved with 1 minute spontaneous activity after each full set at medium luminance (grey screen), which is common practice to estimate ongoing activity^40–42^. Throughout this manuscript, this stimulus type is termed mG, and its associated interleaved grey screen S1. The next set comprised 4 repetitions of 10 temporal frequencies at a SF of 0.03 cpd (stimulus-ON time 14 s, pre- and post-stimulus time 1 s each at grey screen). The stimulus type is referred to as TF and S3. Finally, we presented 15 repetitions of a 60 s grey-scale movie consisting of 2 continuous presentations of the same 30 s movie showing natural scenes such as grass and trees. Each 60 s of movie presentation was followed by a 1 minute grey screen, as a proxy for SA. Movie stimuli are referred to as nat and its SA as S2. Fig. 1 (A, B) outline the stimulus type presentation structure. *Stimulus type* or *stimulus condition* refer to the sets of electrophysiological recordings involving mG, S1, nat, S2, TF and S3. The mouse was placed 25 cm away from the monitor (Samsung SyncMaster 2233Z, 22” LCD monitor, 60 Hz refresh rate, reported as particularly suitable because of its temporal reliability for visual research^43^), covering approximately 60°x75° of the visual field of the right eye. The stimuli on the gamma-corrected monitor were displayed on 255 grey scale with mean screen luminance at 46.93 cd/m2. The left eye was treated with eye ointment (Allergan Lacrilube) and covered with black tape to avoid confounding effects attributable to the binocular zone of vision or inputs from the contralateral eye. The screen output signal and stimulus presentation triggers were synchronised with a custom-built photo-diode circuit board and photo-sensor (LCM555CN) attached to the bottom left corner of the monitor. Here, on stimulus onset and offset a small rectangle flashed, which was detected by the photo-diode, relayed to the data acquisition system and used as a sync pulse.

Stimuli were generated with FlyMouse, a software based on FlyFly, a Matlab Psychophysics Toolbox-based interface developed by the Motion Vision Group at Uppsala University (http://www.flyfly.se/about.html), customized by Silvia Ardila Jimenéz and Marie Tolkiehn. Examples of the DIR and SF stimulus battery are depicted in Fig. 1 (C). SFs were [0.01, 0.02, 0.04, 0.08, 0.16, 0.32] cpd and directions [0°, 45°, 90°, 135°, 180°, 225°, 270°, 325°]. Temporal frequencies were [0.2, 0.4, 0. 6, 1.2, 1.6, 2.4, 3.2, 4.8, 6.4, 9.6] Hz. Signals were acquired by Ripple Grapevine (Scout Processor), amplified with a single-reference amplifier with on-board filtering and digitization at 16 bit resolution and 0.2 μ V/bit (Grapevine Nano front end), and software Trellis, which was equipped with a live display of the channels during the recording. To ensure safe signal transmission, this set-up was best used with an Intel GIGABIT CT DESKTOP RJ45 PCIE B networks card. Broad-band signals sampled at 30 kHz were recorded, filtered between 0.3 Hz and 7.5 kHz (3rd order Butterworth filter) from each of the channels. The electrophysiological data was high-pass filtered, and thresholded at 4 standard deviations, in a spike detection algorithm developed by Aman Saleem (unpublished).

### Analysis methods

Statistical analyses were conducted using MATLAB. The type of the statistical analysis conducted for each analysis was described in the manuscript. Statistical significance was set at P<0.05.

Information theory describes the mathematical study of coding of information, and is, thus, a suitable field to quantitatively examine theories how the brain encodes and processes stimulus information. Shannon entropy^44–46^ is an information-theoretic measure quantifying the amount of uncertainty in a signal. It is minimal in deterministic systems and maximal if each possible state is equiprobable. It may vary under different stimulus conditions informing about pattern diversity. It is an often used metric in decoding^47,48^ and was successfully used in quantifying information in retinal ganglion cells^49^. Similarly, comparing information content in terms of MI can cast light on how a neural population carries information about different types of stimuli or other properties^50^.

#### Spatial patterns

Analysis was based on the binary firing vectors at one time bin, here denoted as patterns. Pattern sizes were 16×1 and 32×1 corresponding to 2 and 4 shanks. 16×1 and 32×1 patterns were created by concatenating adjacent 8×1 shanks (given the probe geometry) of each mouse. For 16×1 patterns, this resulted in 24 non-overlapping experimental sets. Entropy was estimated on 16×1 binary patterns. A subset of the channels was used because for entropy estimates of 2^32^ possible states to be reliable would require many more samples. In addition to patterns, population FRs, i.e. the spatial sum across channels, were investigated to probe population activity without its spatial component.

#### Mutual Information

MI was estimated as *I*(*S,R*) = *H*(*R*) – *H*(*R|S*) = *H*(*S*) – *H*(*S|R*), where *H*(*R|S*) is the (sample-size weighted) sum of entropies conditioned on each stimulus. *H*(*R|S*) is called conditional entropy, and can be interpreted as the mean uncertainty about *R* after observing a second random variable S. *H*(*R*) and *H*(*S*) refer to the entropy of response *R* and stimulus *S*, respectively.

#### Divergence measures

Differences in ensemble pattern distributions under different conditions may indicate that varying neural ensembles are involved in generating them, or that the underlying statistics differed (e.g. cortical state changes), implying different encoding strategies. A common way to measure the distance between probability distributions is the KLD^51^, a measure closely related to MI^45,52,53^. KLD is often used in variational learning and approaches that seek to minimize the distance between a model and a data distribution and describes the amount of information lost when a model distribution is used to approximate the true distribution^54–56^ It is a non-negative, *non-symmetric* measure that does not obey the triangle inequality and is zero only if the two distributions are equal^46,52^. Thus, it is not a real distance. In contrast, we use the JSD, which is based on the KLD with the difference in that it is a real metric: It is symmetric and obeys the triangle inequality, and thus, unlike the KLD it measures an actual distance^57^. This renders it a preferable alternative to the KLD, suitable for comparing probability distributions letting us measure the similarity or distance between them^45,46,53,58^. It can be calculated as the average KLD between a mixture distribution and the two distributions *P* and *Q*. Alternatively, it can be calculated as *the entropy of the mixture distribution minus the mixture of the entropies*: 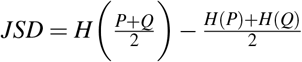 For two probability distributions, and provided using the base-2 logarithm in the entropy estimators, the JSD is bounded between 0 and 1:0 ≤ *JSD* ≤ 1. Thus, although the KLD is often used in the literature, the JSD has its advantages in comparing distributions since it is a real metric. Here, we used pairwise JSD to quantify differences in probability distributions, and as our distance matrix for MDS to visualise the probability distributions distances. MDS was implemented as the *mdscale* function in Matlab (2017a) with default parameters.

## Ethics

The animals were treated in accordance with the Animals Scientific Procedures Act 1986 (United Kingdom) and the Home Office (UK) Animal Care Guidelines. The experiments were approved by the Imperial College Animal Welfare and Ethical Review Board under Project Licence 70/7355 and personal licences.

## Data, code and materials

Shannon entropy was estimated with the Centred Dirichlet Mixture, CDM entropy estimator (for binary discrete data) available at http://github.com/pillowlab/CDMentropy and PYM for discrete data, available at http://github.com/pillowlab/PYMentropy. Data and code of this study are available from the corresponding author upon reasonable request.

## Acknowledgements

We thank Dr. S Mitolo for drawing the mouse in Fig. 1 (A), Dr. S Ardila Jimenéz for useful discussions, and Dr. J Sollini for useful feedback on the manuscript. This research was supported by the Biotechnology and Biological Sciences Research Council (BBSRC grant BB/K001817/1), and a Royal Society Industry Fellowship to SRS. MT was funded by an Imperial College Department of Bioengineering PhD scholarship.

## Author contributions statement

MT carried out the experimental work, data analysis, designed the study, drafted the manuscript, created the figures, performed the statistical analyses; SRS participated in the design of the study and reviewed the manuscript. All authors gave final approval for publication.

## Additional information

The authors declare no competing interests.

